# Rampant loss of social traits during domestication of a *Bacillus subtilis* natural isolate

**DOI:** 10.1101/751438

**Authors:** Hugo C. Barreto, Tiago N. Cordeiro, Adriano O. Henriques, Isabel Gordo

## Abstract

Most well-studied bacteria have been domesticated to some extent. How fast can a natural isolate diverge from its ancestral phenotypes under domestication to a novel laboratory environment is poorly known. Yet such information is key to understand rates of evolution, the time scale at which a natural isolate can be propagated without loss of its natural adaptive traits and the reliability of experimental results across labs. Using experimental evolution, phenotypic assays and whole-genome sequencing, we show that within a week of propagation in a common laboratory environment, a natural isolate of *Bacillus subtilis* acquires mutations that cause changes in a multitude of traits. A single adaptive mutational step, in the gene coding for the transcriptional regulator DegU, impairs a DegU-dependent positive autoregulatory loop and leads to loss of robust biofilm architecture, impaired swarming motility, reduced secretion of exoproteases and changes in the dynamics of sporulation across environments. Importantly, domestication also resulted in improved survival when the bacteria face pressure from cells of the innate immune system. These results show that *degU* is a key target for mutations during domestication and also underscore the importance of performing careful and extremely short-term propagations of natural isolates to conserve the traits encoded in their original genomes.

**Summary:** Domestication is the process by which organisms are selected to live in specific conditions and an important phenomenon that shapes the evolution and variation in many animals and plants. In microbes, domestication is also a key driver of adaptation. It can be beneficial, when improving microbes abilities that are important for biotechnology, but also problematic, especially when studying microbe-host interactions and the microbe’s natural behavior. Using a natural isolate of *Bacillus subtilis*, we determined the speed and genetic basis of microbial domestication using experimental evolution. Within one week of growth in the common laboratory media, mutations in the pleiotropic transcriptional regulator, DegU, emerge and spread in the populations. These lead to loss of social traits, increased resistance to bacteriophages and increased survival in the presence of macrophages. The data highlights the extreme caution that is needed when culturing natural microbial isolates and may help explain why some key microbial social traits and behaviors may differ between different laboratories, even when studying the same strains.

## Introduction

Most bacteria grown in the laboratory face environmental conditions that are distinct from those in their natural habitat. In its natural environment, hardly anywhere a bacterium would find a niche with plentiful nutrients and optimal aeration. Therefore, when sampled from the diverse natural world and subsequently cultured in the lab, bacteria can rapidly adapt and modify its original phenotypes. Such evolutionary domestication can result in increased fitness in the lab at the cost of loss of previous adaptations (1). Some studies in *Escherichia coli*, *Bacillus subtilis*, *Caulobacter crescentus,* and *Saccharomyces cerevisiae* have shown that adaptation to laboratory environments occurs via diverse genotypic paths, but some evolutionary parallelism has also been observed (2–5). The tempo and mode of evolution under domestication is still poorly known. Yet, it is important to quantify how many of the current model bacteria have taken undetermined domestication paths.

The model organism for spore-forming bacteria, *B. subtilis* strain 168, is known to have lost a plasmid and acquired mutations in *sfp*, *epsC*, *swrA*, and *degQ* during its laboratory life (6). These lead to loss of traits likely to be important for *B. subtilis* natural life cycle in the soil, root plants or the gastrointestinal tract of various organisms. Robust biofilm formation is one of such traits, but interestingly, some 168 strains and derivatives are still able to form complex biofilms (7). In addition, a study using genetic engineering introducing the mutated genes and the plasmid showed that it was possible to restore biofilm phenotypes exhibited by the parental less domesticated strain NCIB 3610 (6). Of note is the fact that the mutation in *degQ* leads to a decreased phosphorylation of DegU (8–10), which is a known regulator of social traits in *B. subtilis* (9, 11–13). The strain NCIB 3610 is commonly used as a model for biofilm development. However, it was shown that this strain carries two mutations, one in *rapP* and the other in *dtd* (14), which lead to impaired biofilm formation (15, 16), rendering it less than an ideal model for *B. subtilis* strains with high capacity to form biofilms.

The ability to produce endospores (spores for simplicity) is another important trait of the life cycle of *B. subtilis*. Spores are highly resistant to external stresses and largely responsible for the widespread dissemination of *B. subtilis*. The production of spores is a response to extreme nutrient depletion and under laboratory conditions it is triggered at the onset of the stationary phase of growth (17). A recently characterized natural isolate of *B. subtilis*, BSP1, starts the process of spore formation during growth, unlike its domesticated relatives, and reaches a higher spore titer (18). This occurs as a result of the main activator of the sporulation process, Spo0A∼P, reaching higher levels per cell and in a larger fraction of the population during exponential growth. Spo0A is activated by phosphorylation by means of a phosphorelay that integrates multiple environmental, cell cycle and nutritional cues (17, 18). The precocious and increased sporulation of BSP1 is due to the lack of two genes coding for Rap phosphatases, able to drain phosphoryl groups from the phosphorelay, through the dephosphorylation of Spo0F, a phosphorelay component (18). In accordance with the relevance of sporulation in *B. subtilis* natural life cycle is the fact that under continuous evolution in laboratory conditions a decrease and loss of capability to sporulate was observed (19–22)

The large number of traits that can potentially be lost during continuous growth of *B. subtilis* in classical laboratory environments and the lack of knowledge regarding its first steps of adaptation makes it imperative to understand when and how the domestication occurs. Experimental evolution offers a powerful methodology to study the dynamics of the repeatability of evolutionary change under the same laboratory conditions and to study domestication (19, 23, 24). When coupled with genome sequencing, it allows a real-time assessment of the speed and genetic basis of adaptation to novel environments, as well as the order at which adaptive mutations fix in evolving populations (21, 25–29).

Here we have used experimental evolution to follow the evolutionary path taken by a natural isolate of *B. subtilis* during domestication to a common laboratory environment. We found that within one week (8 passages), mutations in *degU*, coding for a global transcriptional regulator (11, 12, 30), spread and cause a rapid change in several social traits of *B. subtilis*, likely to be important for its fitness in the wild (31). Using gene-editing we show that one of the *degU* mutations causes attenuation of swarming motility, reduction of biofilm production and alteration of its architecture, increased resistance to bacteriophage SPP1 infection and reduced exoprotease secretion. Importantly, the initial process of domestication also resulted in a change of the sporulation dynamics across environments and an increased survival when the bacteria face pressure from the innate immune system. DegU thus emerges as a central mutational target during domestication. Overall, our results indicate that the propagation of natural isolates in the laboratory should be performed with extreme care as domestication can lead to rampant loss of traits that while important for the *B. subtilis* natural life cycle are largely dispensable under laboratory conditions.

## Results

### Emergence of a new adaptive colony morphology during *B. subtilis* domestication.

Five populations derived from a *B. subtilis* natural isolate (BSP1) hereinafter named Ancestral, were daily passaged for sixteen days, through dilution in a rich medium with agitation and aeration. Samples were frozen every two days so that evolutionary steps during short-term domestication could be followed through whole-genome sequencing. Daily plating revealed that, within the first week of the experiment, two new colony morphotypes emerged (Fig. 1A). The ancestral type *a* dominated the initial populations, but a new type *b*, characterized by a flat colony morphotype, and a type *c*, an intermediate morphotype, reached appreciable frequencies rapidly. Type *b* was detected in all the populations, while type *c* was only observed in three out of five populations (Fig. 1B). As type *b* achieved the highest frequency, reaching fixation after sixteen days in population 1 (Fig. 1C), we conducted a detailed phenotypic and genetic characterization of one clone from this population. The rapid spread of type *b*, as well as its repeated emergence, suggests that it carries an advantage when growing in the laboratory environment. To test this hypothesis, we selected a type *b* colony from population 1 at day eight, hereinafter named Evolved, and characterized its growth in LB. Indeed both the maximum growth rate per hour and the carrying capacity after 7 hours of growth of the Evolved were significantly higher than those of the Ancestral (Fig. S1A and B). This shows that Evolved has increased growth traits in the laboratory environment.

**FIG 1.**
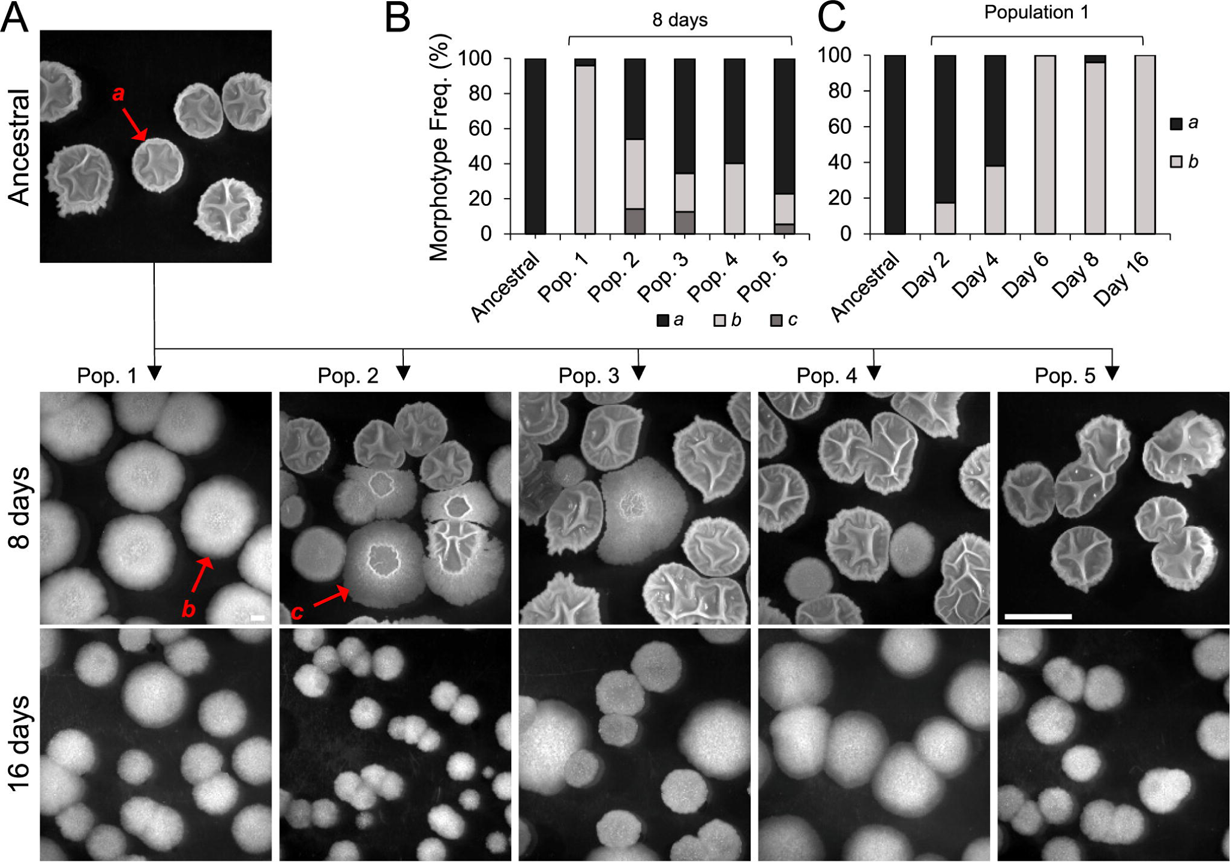
Changes in colony morphology with domestication. (A) Representative image of the Ancestral colony morphology and of the three different types of colony morphology, *a*, *b* and *c*, observed at the eight and sixteen day of the domestication experiment in the five evolved populations; (B) Frequency of each morphotype in the five populations at day 8; (C) Frequency of morphotype *b* in population 1 along time. The scale bar represents 1 cm and applies to all panels.

### DegU as a main target for the adaptation to the laboratory environment

To determine the genetic basis of the adaptive morphotype we performed whole-genome sequencing of the Ancestral and the Evolved clone. Only one mutation was observed, in the *degU* gene, coding for the response regulator DegU (32). The non-synonymous mutation is a T-to-G transition causing the substitution of isoleucine 186, in the helix-turn-helix motif (HTH), within the DNA-binding domain of DegU, by a methionine (I186M) (32) (Fig. 2A). *degU* is part of the *degS*-*degU* operon, coding for a two-component system that controls social behaviors in *B. subtilis* (9, 11–13). Interestingly, the laboratory strain 168 is known to have a mutation in *degQ* that leads to a decreased phosphorylation of DegU (8–10). This suggested that the I186M substitution could cause an adaptive change capable of affecting several traits. In the laboratory strain of *B. subtilis* the unphosphorylated form of DegU activates genes involved in the ability to uptake external DNA during competence development (33). During growth, the intracellular concentration of the phosphorylated form of DegU (DegU∼P), increases, leading to the progressive activation of genes required for swarming motility, biofilm formation and the secretion of extracellular enzymes (9, 34). Given the central importance of DegU in processes that could be costly in the laboratory environment we tested whether additional clones of the natural isolate BSP1, that had evolved independently, also carried mutations in *degU*. Sanger sequencing revealed that all the five clones isolated from each population had mutations in *degU*: in addition to the mutation causing the I186M substitution (Evolved, in population 1; see above), the same mutation was identified in a clone from population 4; a mutation causing the substitution of histidine 200 by a tyrosine (H200Y) was identified in two populations (in population 1 and in population 5), and another, causing the replacement of valine 131 by an aspartate (V131D) was identified in one (population 2) (Fig. 2A and B). Finally, one mutation caused the insertion of a TGA codon leading to premature translation arrest after codon D18 (Fig. 2A and C; population 3). These populations exhibited distinct colony morphologies but all appeared less structured than the colonies formed by Ancestral (Fig. 2C). The V131D substitution lies within the linker region that separates the receiver and DNA-binding domains of DegU, while H200Y (as for I186M; above) is located in the HTH motif (Fig. 2B). All of the missense mutations affect amino acid residues that are conserved among DegU orthologs (Fig. S2A and B) and are thus likely to be functionally important. In particular, the I186M and H200Y substitutions are likely to affect DNA binding (Fig. 2A; see also Text S1 for a discussion of the effect of the various substitutions). These results suggest a high degree of evolutionary parallelism at the gene level and implicate DegU as the first main target of domestication to the laboratory for the natural isolate BSP1.

**FIG 2.**
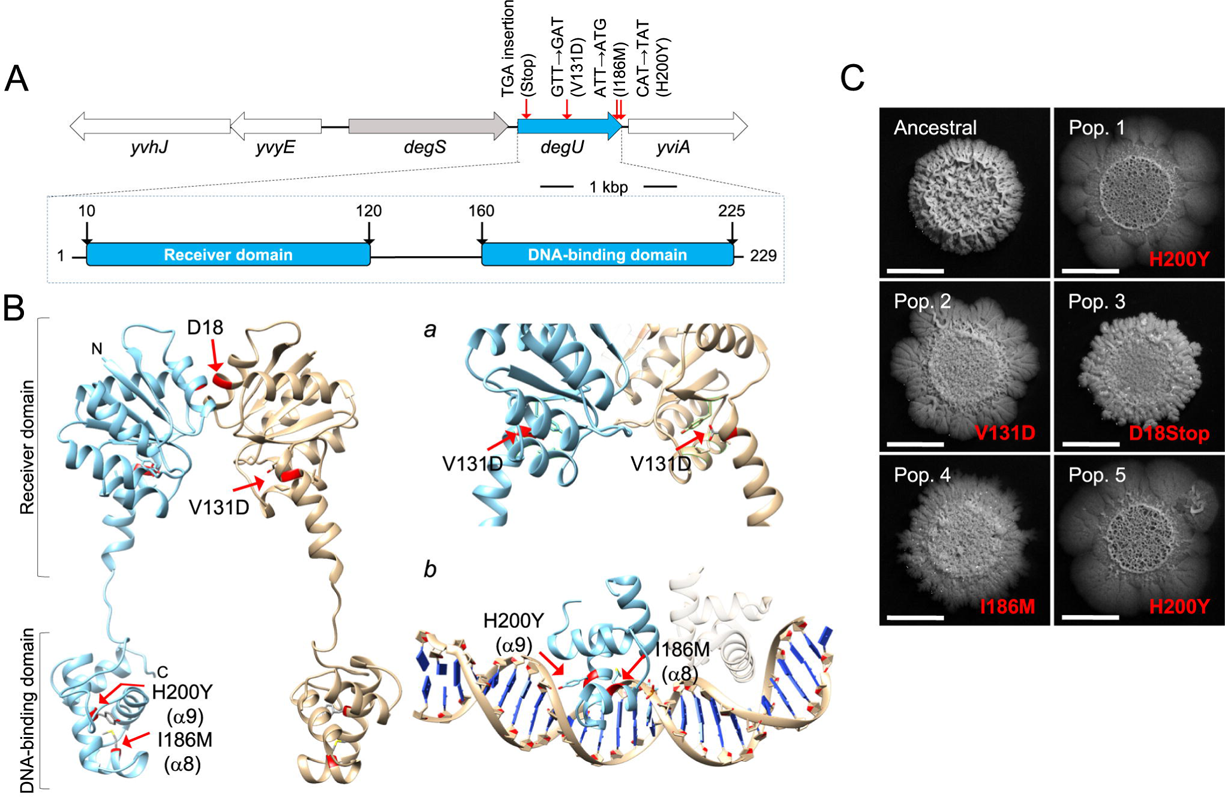
Domestication is accompanied by mutations in *degU.* (A) *degU* region of the *B. subtilis* chromosome (top) and domain organization of the DegU protein (bottom). The position of the various mutations found and the corresponding amino acid substitution is indicated. (B) Model of the full-length DegU protein *of B. subtilis* obtained by comparative modeling and using the crystal structure of the LiaR protein from *Enterococcus faecalis* as the template (PDB code: 5hev). The protein is thought to form a dimer and the two monomers are represented in blue and light brown, with the position of the receiver and DNA-binding domains indicated. The red arrows indicate the location of the single amino acid substitutions found in DegU. *a* and *b*, show a magnification of the regions encompassing the V131D (*a*) and I186M and H200Y (*b*) substitutions. In *b*, the region of the helix-turn-helix (HTH) motif is modeled with DNA, to highlight the likely involvement of residues I186 and H200 in DNA binding. The HTH motif was independently modeled using the crystal structure of the LiaR DNA-binding domain as the template (PDB code: 4wuh). (C) Representative images showing the complex biofilm morphology of Ancestral and clones representative of each population after 16 days of domestication. In red are indicated the mutation in DegU present in each clone. All strains were incubated in MSgg for 96h at 28°C. Scale bar, 1 cm.

### The I186M substitution in DegU is responsible for the colony phenotype of Evolved

We proceeded with a detailed characterization of the Evolved from population 1 which carried a I186M substitution in DegU. To test whether this mutation caused the alteration in colony morphology, the wildtype *degU* allele (or *degU^Anc^*, for Ancestral) and the allele coding for DegU^I186M^ (found in population 1, or *degU^Evo^*) were introduced ectopically at the non-essential *amyE* gene in a strain bearing a *degU* knockout constructed in the background of Ancestral (Fig. S3; see also Text S1). The resulting strains, termed *degU^Anc^* and *degU^Evo^*, had a colony morphology indistinguishable from Ancestral and Evolved / *degU^Evo^*respectively (Fig. 3C). We infer that the *degU^Evo^* allele is responsible for the colony phenotype of Evolved.

**FIG 3.**
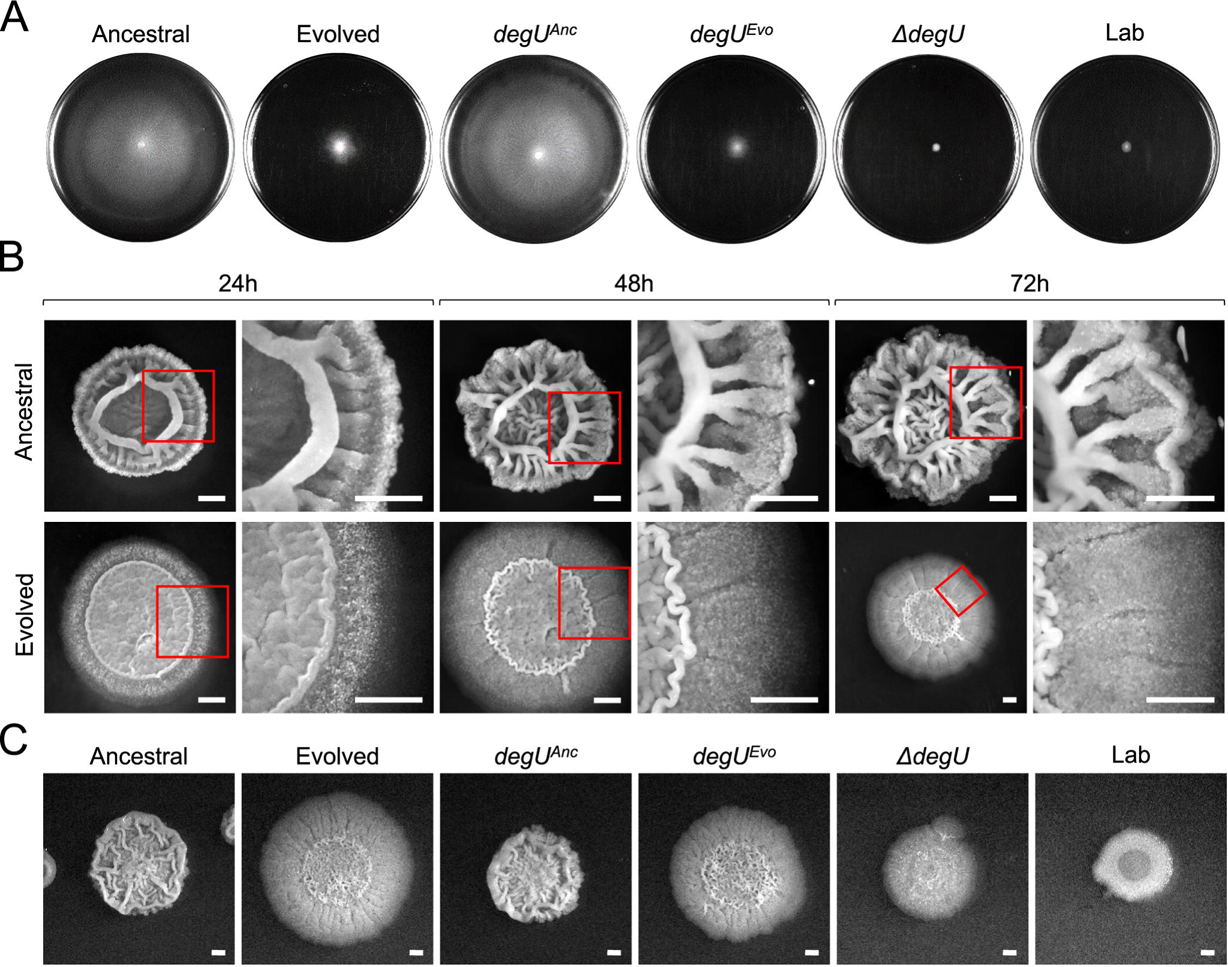
*degU^Evo^* is responsible for the alteration in swarming motility and colony architecture. (A) Swarming motility assay of Ancestral, Evolved, *degU^Anc^*, *degU^Evo^*, Δ*degU*, and Lab. LB plates fortified with 0.7% of agar were inoculated incubated for 16h at 28°C. Swarm expansion, resulting from bacterial growth, appears in white, whereas uncolonized agar appears in black. (B) Representative images showing the complex colony architecture development along with the indicated time points of the Ancestral and Evolved. The strains were grown in MSgg medium at 28°C. (C) Representative images showing the complex colony architecture of the indicated strains on MSgg agar plates incubated for 96h at 28°C. Scale bars, 1 cm.

### *degU*^I186M^ causes adaptive loss of major social traits

In commonly used laboratory strains of *B. subtilis*, such as 168 and NCIB 3610, several social traits are regulated by the unphosphorylated form as well as by low, medium or high levels of DegU∼P (12, 30). DegU functions as a “rheostat”, sensing environmental signals and allowing the expression of competence, social motility (or swarming), biofilm development and exoprotease production along a gradient in the cellular accumulation of Deg∼P (12). Competence development is positively regulated by the unphosphorylated form of DegU (33, 35, 36). ComK is a regulatory protein required for competence development that drives transcription of the genes coding for the DNA uptake and integration machinery but also stimulates transcription of its own gene (37). Unphosphorylated DegU functions as a priming protein in competence development by binding to the *comK* promoter and facilitating ComK stimulation of *comK* transcription at low ComK concentrations (33, 37, 38). Given this, we proceeded to test if these phenotypes observed in laboratory strains were also regulated by DegU in the natural isolate strain BSP1, as well as the effect of the I186M substitution in the Evolved. We found no differences in the development of competence between Ancestral or *degU^Anc^* and Evolved or *degU^Evo^* (Fig. S4). This suggests that the I186M substitution does not affect the function of the unphosphorylated form of DegU in promoting competence development and thus that *degU^Anc^* is not a loss-of-function allele. Low levels of DegU∼P, however, activate transcription of genes involved in social motility, or swarming (9, 11, 39, 40). Swarming motility assays revealed that while the Ancestral and *degU^Anc^* have the ability to swarm, the Evolved and *degU^Evo^* show poor swarming ability (Fig. 3A). A widely used laboratory strain, PY79, hereinafter termed Lab, as well as other laboratory strains, carry mutations that prevent swarming motility (41–43). In our assay, Lab as well as a *degU* insertional mutant, are deficient in swarming motility (Fig. 3A). Thus, the I186M substitution leads to limited pleiotropic effects, *i.e*., decreased swarming motility without affecting competence.

The production of proteins responsible for biofilm formation is another trait under the control of DegU∼P (8, 9, 11, 44, 45). To query whether biofilm architecture and robustness were affected by the I186M substitution, we examined both the colony architecture over time, the colony being a biofilm formed at the solid medium/air interface, as well as the formation of biofilms at the liquid/air interface in liquid cultures. Both Ancestral and *degU^Anc^*showed a complex colony architecture characterized by many wrinkles after 24 h of incubation, and the complexity of the colony architecture increased with time during the assay (Fig. 3B). In contrast, Evolved and *degU^Evo^* formed colonies that tended to be flatter and with fewer wrinkles (Fig. 3B and C). As a control, colonies formed by a *degU* insertional mutant show extremely low complexity and compared to Ancestral, Lab also formed colonies with a simpler architecture (Fig. 3C). These results are consistent with the importance of DegU for the formation of a biofilm at a solid medium/air interface (44, 45). Similarly, in liquid cultures, both Evolved and *degU^Evo^* formed a biofilm at the liquid/air interface less robust than that of the Ancestral as determined by the quantification of the pellicle formed (Fig. 4A and Fig. S5). Thus, *degU^Evo^* affects the ability of *B. subtilis* to form complex, robust biofilms.

**FIG 4.**
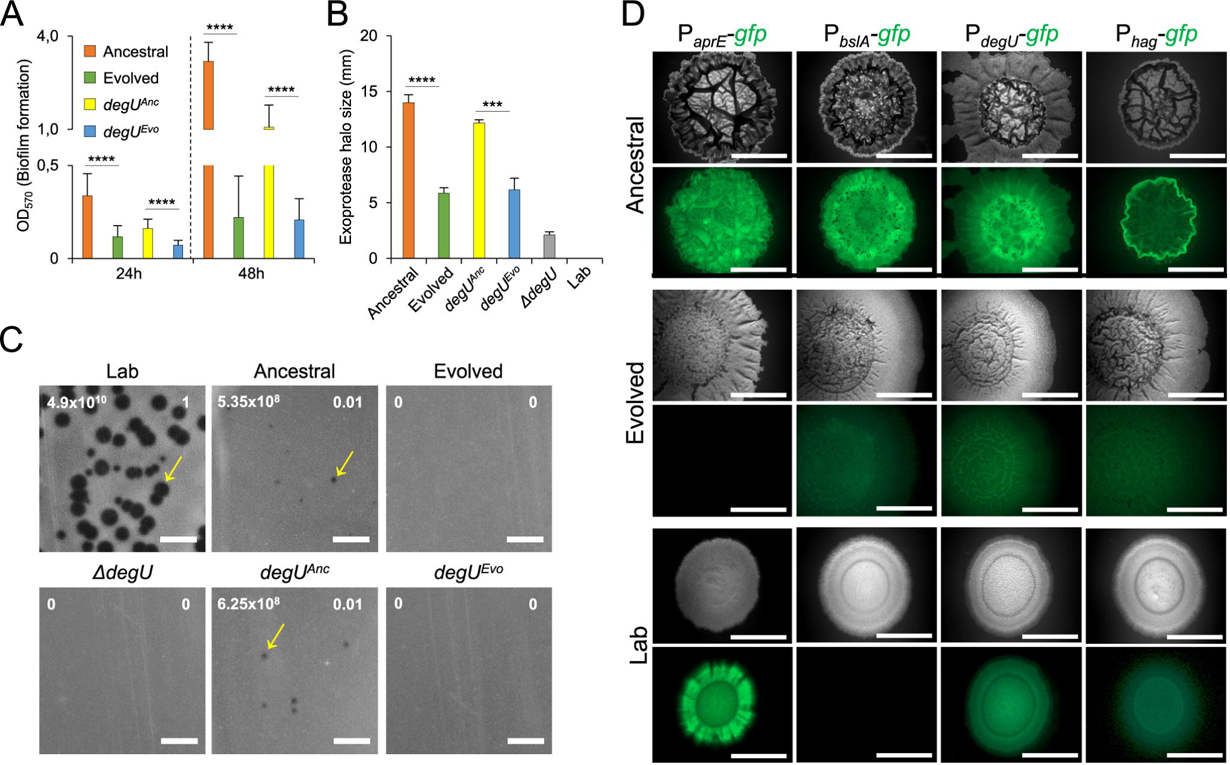
*degU^Evo^* is responsible for the alteration in biofilm complexity, exoprotease secretion, phage resistance and the pattern of gene expression during biofilm formation. (A) Quantification by the crystal violet method of the biofilms formed by the Ancestral (n = 18 for 24h, n = 15 for 48h), Evolved (n = 22 for 24h, n = 15 for 48h), *degU^Anc^* (n = 18 for 24h, n = 19 for 48h), and *degU^Evo^* (n = 19 for 24h, n = 18 for 48h) in MSgg broth incubated at 25°C for the indicated time points. For the statistics the Mann-Whitney U test was used. For ****, p<0.0001. (B) Dimension of the halos produced by the Ancestral (n = 4), Evolved (n = 4), *degU^Anc^* (n = 3), *degU^Evo^* (n = 3), and Δ*degU* (n = 4) in LB fortified with 1.5 % agar and supplemented with 2% of skimmed milk incubated at 37°C for 48h. For the statistics an unpaired t-test with Welch’s correction was used. For ****, p<0.0001, for ***, p=0.007. The error bar represents the standard deviation. (C) Efficiency of Plating (EOP) shown in white numbers in the superior right corner for the Ancestral, Evolved, *degU^Anc^*, *degU^Evo^*, Δ*degU* using as a reference the indicator strain Lab (PY79), which is phage sensitive. The number of plaque forming units (PFU’s) is shown in the superior left corner in white numbers. The yellow arrows indicate SPP1 phage plaques. Note that Ancestral is sensitive to SPP1 but the plaque size is reduced when compared with the Lab strain, while the Evolved is resistant. Scale bars, 1 cm. (D) Representative images of the expression of transcriptional fusions between the *aprE*, *bslA*, *hag* and *degU* promoter regions and *gfp* in Ancestral, Evolved, and Lab after 96h of incubation in MSgg at 28°C. Scale bar, 1 cm. In panel A and B the error bars represent the standard deviation.

Other processes important for *B. subitlis* social natural lifestyle that depend on high levels of DegU∼P include the secretion of exoproteases and survival under bacteriophage predation (12, 39, 46, 47). We found decreased exoprotease secretion in Evolved and the *degU^Evo^* strain, as compared to Ancestral and *degU^Anc^*(Fig. 4B). For reference, exoprotease secretion was severely impaired in both the *degU* insertional mutant and in Lab (Fig. 4B). Lastly, when infected with the bacteriophage SPP1, both Evolved and *degU^Evo^* showed increased resistance to phage infection, as compared to Ancestral and *degU^Anc^* or Lab (Fig. 4C). Overall, these results show that in the natural isolate BSP1, as in the laboratory strain, DegU is a key regulator of social straits. They also reinforce the view that the *degU^Evo^* mutation is pleiotropic but not fully, as it affects the phenotypes regulated by DegU∼P, including social mobility, biofilm formation, exoprotease production and resistance to phage infection, but does not impair the function of unphosphorylated DegU in priming competence.

### Decreased transcription of DegU∼P-target genes after domestication

The phenotypic assays performed showed that in the natural isolate BSP1 the I186M substitution in DegU changed traits regulated by DegU∼P in the first steps of domestication. The location of the I186M substitution in the HTH motif of DegU raised the possibility that the transcription of genes regulated by DegU∼P could be impaired in Evolved in comparison with Ancestral. To test this, we constructed transcriptional fusions between the promoters for the *hag*, *bslA*, and *aprE* genes and the *gfp* gene. *hag* codes for flagellin, the main component of the flagellum and is required for social motility; and the expression of *hag* indirectly requires low levels of DegU∼P (11, 48). *bslA* codes for a self-assembling hydrophobin that forms a hydrophobic coat at the surface of biofilms; as shown in strain NC3610, a model for biofilm development, BslA is required for the formation of structurally complex colonies and biofilms (49–52). Lastly, *aprE* codes for subtilisin, a major alkaline exoprotease (32, 53). Transcription of both the *bslA* and *aprE* genes is subject to a logic AND gate, in that it requires both derepression of both promoters under the control of Spo0A∼P and in addition, DegU∼P (11, 32, 49, 54).

We then examined the transcription of these genes at the population (biofilms) and single-cell levels. In Ancestral, the P*_aprE_*-, P*_bslA_*- and P*_degU_*-*gfp* fusions were expressed throughout the architecturally complex colonies, whereas expression of P*_hag_*-*gfp* was expressed mostly at the colony edge (Fig. 4D). Expression of P*_aprE_*-, P*_degU_*- and P*_hag_*-*gfp* was also detected in Lab, but at lower levels, and expression of P*_bslA_-gfp* was not detected, consistent with the simpler colony morphology (Fig. 4D). Interestingly, in spite of the maintenance of a complex colony architecture, expression of P*_bslA_*-, P*_degU_*- and P*_hag_*-*gfp* was detected at very low levels in Evolved, and expression of P*_aprE_*-*gfp* was not detected (Fig. 4D). Thus, DegU^I186M^ reduces the expression of DegU target genes during biofilm development.

During planktonic growth, transcription of *hag* decreases markedly in the absence of DegU∼P (9). Moreover, expression of *hag* is heterogeneous, with free cells showing higher expression levels than chained cells (55). Accordingly, in Ancestral, the free cells showed higher fluorescence intensity from the P*_hag_*-*gfp* fusion than chained cells (p < 0.0001, Kruskal-Wallis test with Dunn’s test of multiple comparisons; Fig. 5A and B). In contrast, Evolved showed a decrease in the GFP signal from P*_hag_*-*gfp* both in free cells (∼2.2 fold; p < 0.0001, Kruskal-Wallis test with Dunn’s test of multiple comparisons) and chains (∼2.8 fold; p < 0.0001, Kruskal-Wallis test with Dunn’s test of multiple comparisons), relative to the Ancestral (Fig. 5B). Although Lab does not exhibit swarming motility, as previously reported (55) it showed increased expression of P*_hag_*-*gfp* in both free cells (p < 0.0001, Kruskal-Wallis test with Dunn’s test of multiple comparisons) and chains (p < 0.0001, Kruskal-Wallis test with Dunn’s test of multiple comparisons) when compared with Ancestral, which exhibits swarming motility (Fig. 5B). Expression of *bslA* is heterogeneous between free cells and chains in the Ancestral, as also found for *hag* (Fig. 5A; see also above), and was ∼2.8 fold lower in free cells when compared to chained cells (p < 0.0001, Kruskal-Wallis test with Dunn’s test of multiple comparisons; Fig. 5B). Strikingly, no heterogeneous expression between free cells and chains of *bslA* was observed in the Evolved clone (Fig. 5B). Moreover, the level of *bslA* expression was similar in free cells and chains (p = 0.0889, Kruskal-Wallis test with Dunn’s test of multiple comparisons), although *bslA* expression was ∼1.5 fold lower (p < 0.0001, Kruskal-Wallis test with Dunn’s test of multiple comparisons) in the free cells when compared to Ancestral (Fig. 5B). In the Lab strain, which does not produce robust biofilms, transcription of *bslA* is markedly reduced (Fig. 5B). In laboratory strains, DegU∼P is a direct positive regulator of *aprE* and expression of *aprE* was shown to be bi-stable in a laboratory strain (54). Consistent with this finding, low and high levels of *aprE* expression were detected for Lab, both in free cells and chains (Fig. 5). In contrast, Ancestral did not express high levels of *aprE*, although it showed heterogeneity between free cells and chains in *aprE* expression (Fig. 5). Lastly, Evolved showed greatly reduced expression of *aprE*, both in free cells (p < 0.0001, Kruskal-Wallis test with Dunn’s test of multiple comparisons) and chains (p < 0.0001, Kruskal-Wallis test with Dunn’s test of multiple comparisons) (Fig. 5). Thus, domestication in the natural isolate BSP1 is accompanied by a sharp decrease in the expression of *aprE*.

**FIG 5.**
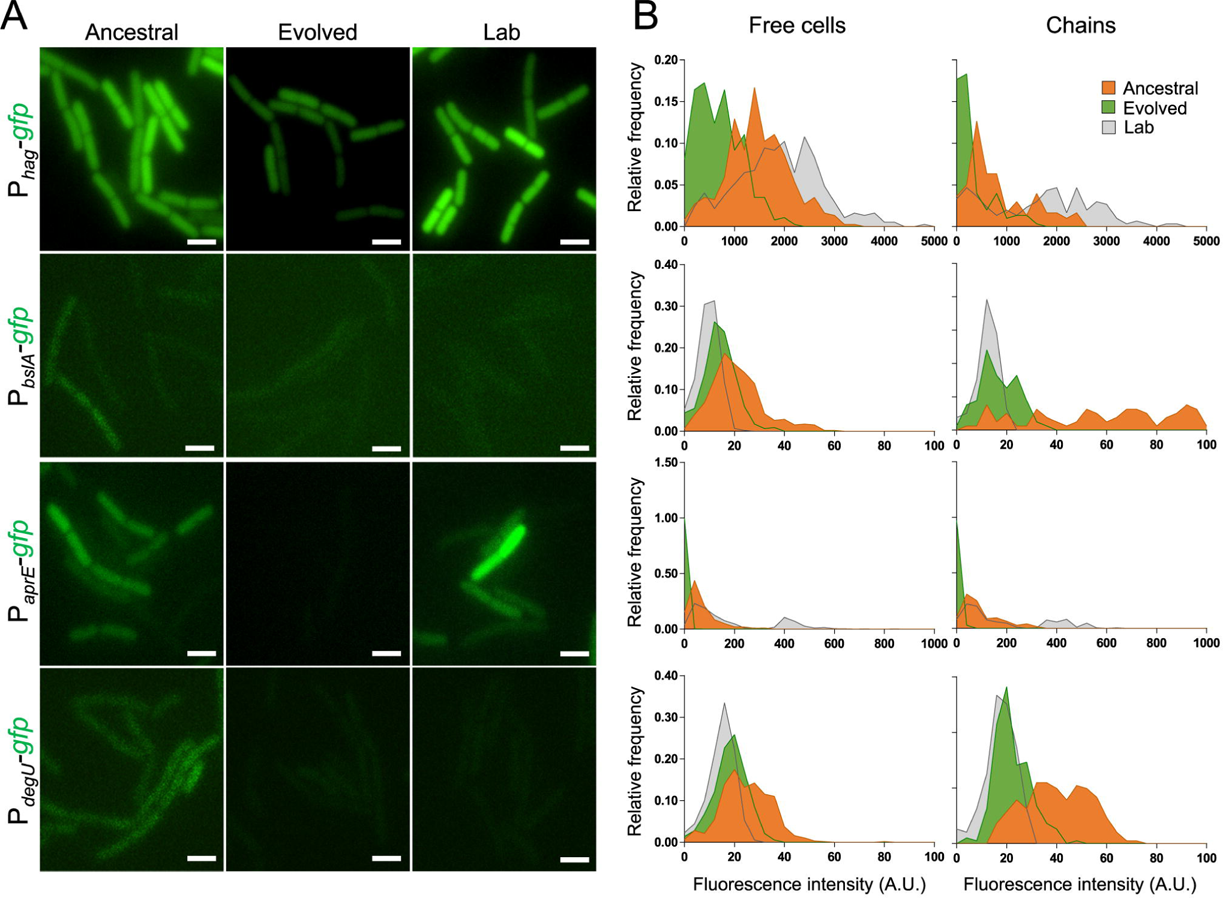
*degU^Evo^* alters the pattern of gene expression at the single-cell level. (A) Representative images of the expression of *hag-*, *bslA-*, *aprE-* and *degU-gfp* transcriptional fusions in Ancestral, Evolved, and Lab one hour after the onset of stationary phase in LB. The cultures were grown with agitation at 37°C. Scale bar, 1 µm. (B) Relative frequency of expression of transcriptional fusions of the indicated promoters to *gfp* in the same conditions as above. For the relative frequency of expression of transcriptional fusions in free cells, a total of 371 (*hag-gfp*), 453 (*bslA-gfp*), 713 (*aprE-gfp*), and 561 (*degU-gfp*) cells from Ancestral, Evolved and Lab were analysed. For the relative frequency of expression of transcriptional fusions in chains, a total of 150 (*hag-gfp*), 79 (*bslA-gfp*), 168 (*aprE-gfp*), and 191 (*degU-gfp*) cells from Ancestral, Evolved and Lab were analysed.

Overall, these results show that in the natural isolate BSP1, the I186M substitution in DegU reduces the expression of DegU∼P regulated genes during biofilm development and planktonic growth.

### Domestication impairs a *degU* positive auto-regulatory loop

In its high-level phosphorylated state, DegU∼P activates transcription of *degU* itself (56, 57) by binding to a site in the *degU* regulatory region (34, 58). Importantly, this positive auto-regulatory loop contributes to the heterogeneous expression of *degU* and the DegU∼P-dependent genes in laboratory strains (54). Since Evolved shows reduced expression of DegU∼P-target genes, we wanted to test whether DegU^I186M^ also impaired expression of *degU*, reducing the levels of DegU. We found expression of P*_degU_-gfp* to be heterogeneous between free cells and chains in Ancestral, with chains showing a ∼1.6 fold higher expression relative to free cells (p < 0.0001, Kruskal-Wallis test with Dunn’s test of multiple comparisons; Fig. 5) (55). In Evolved and Lab strains, expression of *degU* was reduced when compared to Ancestral, both in free cells (p < 0.0001, Kruskal-Wallis test with Dunn’s test of multiple comparisons) and chains (p < 0.0001, Kruskal-Wallis test with Dunn’s test of multiple comparisons) (Fig. 5).

Together, these results suggest that I186M impairs the ability of DegU∼P to activate transcription of *degU* itself. Impaired activation of the DegU auto-regulatory loop, in turn, reduces transcription of degU itself, suggesting an explanation to why the expression of genes directly or indirectly regulated by DegU∼P in Evolved is reduced.

### Domestication leads to increased survival in the presence of cells of the immune system

*B. subtilis* has been isolated from the gastrointestinal tract of several animals including humans (59, 60) and has been found to grow, sporulate and persist in the murine gut (61, 62). Thus, it can experience selective pressures inside a host. To determine if domestication could impair the ability of *B. subtilis* to withstand a hostile host environment, we measured the survival of Evolved and Ancestral in the presence of cells of the innate immune system - macrophages. Interestingly, the Evolved strain showed an increased survival over the Ancestral in the presence of macrophages, both at 3h (p = 0.001) and 5h (p < 0.0001). (Fig. 6A). This result shows that the domestication to the laboratory environment coincidently leads to a survival advantage for *B. subtilis* when facing cells of the host immune system.

**FIG 6.**
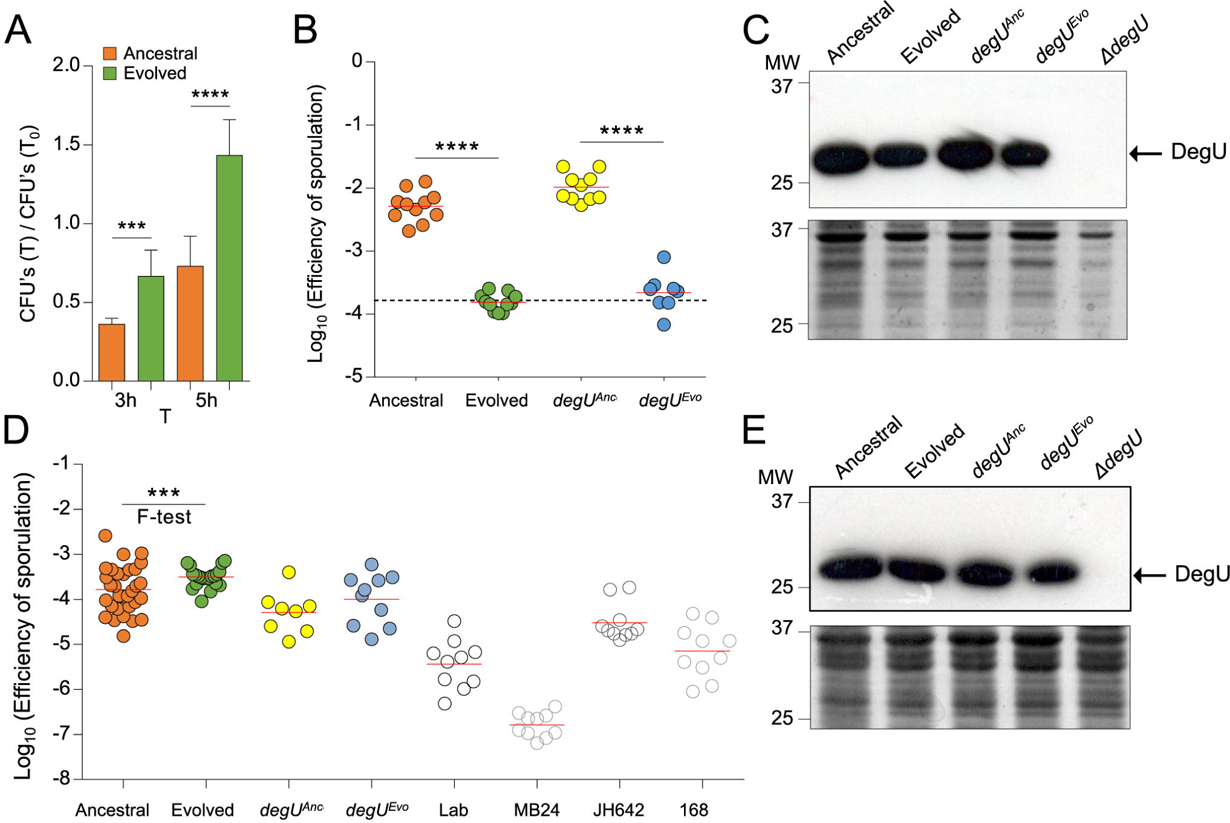
degU^Evo^ increases survival in the presence of macrophages and changes sporulation efficiency in an environment-dependent manner. (A) Macrophages were infected with Ancestral (n = 9) and Evolved (n = 9) and colony-forming units of both the intracellular and extracellular bacteria obtained by plating at the indicated time points. For the statistics the Unpaired t test with Welch’s correction was used. ***p = 0.001, ****p < 0.0001 The error bar represents the standard deviation. (B) Comparison of the sporulation efficiency in RPMI medium between Ancestral (n = 11), Evolved (n = 11), *degU^Anc^* (n = 10) and *degU^Evo^*(n = 8). The efficiency of sporulation was calculated as the ratio between the heat resistant spore counts and total (viable) cells. The dashed line indicates the average sporulation efficiency for the Ancestral in LB. For the statistics an ANOVA and Tukey’s multiple comparison test was used. For ****, p<0.0001. (C) Accumulation of DegU in Ancestral, Evolved, *degU^Anc^*, *degU^Evo^* and the *degU* insertional mutant in RPMI. (D) Comparison of the sporulation efficiency and variance in LB between Ancestral (n = 31), Evolved (n = 21), *degU^Anc^* (n = 8), *degU^Evo^* (n = 10), Lab (n = 10) and three other commonly used laboratory strains (MB24, n = 10, JH642, n = 10, and 168, n = 10). For the mean sporulation efficiency an ANOVA and Tukey’s multiple comparison test was used. For the variance the F-test was used. For ***, p=0.0002. (E) The levels of DegU are similar between Ancestral and Evolved in LB. Accumulation of DegU in Ancestral, Evolved, *degU^Anc^*, *degU^Evo^*, and the *degU* insertional mutant. In (C) and (E), the cells were collected after growth in RPMI (C) or LB (E) and whole-cell lysates prepared (see Methods). Proteins (20 µg) in whole-cell lysates were resolved by SDS-PAGE and subject to immunoblot analysis with an anti-DegU antibody. The arrow shows the position of DegU; the red arrows indicate slightly higher levels of DegU. The panel below the immunoblot shows part of a Coomassie-stained gel, run in parallel, as a loading control. The position of molecular weight markers (in kDa) is shown on the left side of the panels. In panels B and D the red line indicates the mean.

### Sporulation efficiency changes across environments

High levels of DegU∼P promote sporulation by increasing the levels of Spo0A∼P, the master regulatory protein for entry into sporulation (13). Since the I186M substitution reduced transcription of *degU*, we reasoned that changes in the frequency of sporulation could also have occurred during domestication. We first tested this phenotype in a host-related environment (RPMI medium) and found the sporulation efficiency of Ancestral (and of *degU^Anc^*) to be ∼1.5 Log10-fold higher than that of Evolved (or *degU^Evo^*) (Fig. 6B). The levels of DegU, as assessed by immunoblot analysis with an anti-DegU antibody of established specificity (63), are slightly higher in RPMI for both Ancestral and *degU^Anc^*as compared to Evolved or *degU^Evo^* (Fig. 6C).

We also tested the ability of our strains to sporulate in the environment where the domestication process occurred (LB medium). Most laboratory strains sporulate at very low levels in LB (about 10^4^ spores/ml of culture as compared to 10^8^ spores/ml in a medium such as DSM that support efficient sporulation) (18). We found no significant difference in the mean efficiency of sporulation between Evolved (and *degU^Evo^*) and Ancestral (or *degU^Anc^*) in LB (Fig. 6D). Interestingly, the variance was reduced in the Evolved when compared with the Ancestral (F-test, p = 0.0002). In addition, all of the laboratory strains tested sporulated in LB at efficiencies lower than that of the Ancestral, although one strain, JH642, sporulated better than the other laboratory strains tested under these conditions (Fig. 6D). This is consistent with the initial description of Ancestral, which enters sporulation during growth and reaches a higher titer of spores than any of the laboratory strains tested in DSM (18). The steady-state levels of DegU in whole-cell lysates obtained 1 hour after the onset of stationary phase in LB revealed very similar levels of DegU for all the strains (Fig. 6E).

These results suggest that the I186M mutation can affect the intracellular levels of DegU in a manner that depends on the environment and show that the influence of DegU on sporulation exhibits antagonistic pleiotropy.

## Discussion

BSP1 is a gastro-intestinal isolate of *B. subtilis* in which sporulation initiates during growth. This happens because BSP1 lacks three *rap* genes, coding for phosphatases that normally drain phosphoryl groups from the phosphorelay, thus negatively regulating the activation of Spo0A. As such, more cells in the population have Spo0A active above a threshold level required to induce sporulation. Several other gastro-intestinal isolates of *B. subtilis*, including from the human gut, also lack combinations of the *rap* genes and show enhanced sporulation (18, 62). *B. subtilis* completes its entire life cycle in the gut (62), and it seems likely that sporulation is important for survival and/or propagation in the gut ecosystem, as shown for other spore-formers (64, 65). Sporulation is also important for the efficient dispersal of spore-formers through the environment and among hosts (66–68). Sporulation is, however, a time and energetically costly process, requiring the differential expression of over 10% of the genome over a period of 7-8 hours (18, 69). Accordingly, the propagation of *B. subtilis* in the laboratory in the absence of selection for sporulation results in a reduction in the ability to sporulate (19–22). *B. subtilis* has been used in laboratory conditions for more than fifty years, and has accumulated mutations likely to be adaptive in that environment and that, relative to wild strains, lead to the attenuation of phenotypic traits that include swarming motility (41, 42), poly-γ-glutamate synthesis (8), production of antibiotics, the secretion of degradative enzymes (70) or the formation of robust biofilms (6, 8, 71). These processes have either become neutral with respect to fitness, or selection favored its loss under laboratory conditions. In addition, repeatadly selecting individual colonies to cultivate and maintain bacteria in laboratory conditions can increase the chances of loss of phenotypes independently of fitness differences (72).

Here we traced one example of possible domestication routes of the natural isolate *B. subtilis* strain BSP1. Rapid changes in colony morphotypes were observed in parallel cultures, leading to complete fixation of a specific colonial morphotype, termed type *b*, in all the replicate cultures after two weeks (Fig. 1). The adaptive morphotype is characterized by a smooth and flat colony, lacking the complex architectural features of the original strain (Fig. 2C). Similar colony morphology changes were previously observed during domestication of other *B. subtilis* strains (3, 19–21, 24). Colonies are biofilms formed at the solid/air interface (1). As such, these observations hinted at attenuation of an important social behavior. Studies in *Salmonella enterica*, *Saccharomyces cerevisiae*, *Bacillus licheniformis, Aneurinibacillus migulanus*, and *Myxococcus zanthus* have also documented the appearance of smooth colonies within a short period of time (5, 73–76). This suggests that phenotypic parallelism across species is a broad pattern of adaptation to the laboratory environment.

At the genomic level we detected mutations in the coding region of the *degU* gene (Fig. 2). DegU is the response regulator of the two-component system DegS-DegU and controls social traits such as biofilm formation, swarming motility and exoprotease secretion (11). In its non-phosphorylated state, DegU is responsible for the development of competence while the rise in DegU∼P levels sequentially activates swarming, biofilm formation and exoprotease secretion (9). DegU belongs to the NarL/FixJ subfamily of DNA binding proteins (77). We characterized in detail the effects of a mutation leading to the I186M substitution in DegU. The I186M substitution occurs in the DNA-recognition helix of the DegU HTH motif, located in the C-terminal domain of the protein (Fig. 2A). Modeling studies indicate that this substitution is likely to affect a contact of the HTH motif with bases in the major groove of DNA as also shown in the crystal structure of NarL, in which I186 is conserved, with DNA (78) (Fig. 2 and Fig. S2; see also Text S1). Moreover, alanine-scanning mutagenesis showed decreased transcription of the DegU-controlled genes *comG* (as a proxy for the activity of ComK, a direct target of DegU) and *aprE* in a strain producing DegU^I186A^, and the substitution also affected the binding of DegU to the *comK* and *aprE* target promoters (32). I186 is also conserved in LuxR, another NarL family member, and its replacement by Ala also resulted in reduced binding to target DNA sequences (79). One other substitution found in DegU in our study, H200Y, is also likely to impair DNA binding, as suggested by the study of a single Ala substitution in transcription and DNA binding to cognate sites in the promoters of the DegU-responsive genes *comK* and *aprE* (32). This residue, however, as suggested by the structure of a NArL-DNA co-crystal, contacts the DNA phosphate backbone (78) (see also S1 Text). Importantly, while the I186M substitution found in the domesticated clone, Evolved, did not cause changes in the efficiency of transformation with exogenous DNA, it impaired processes regulated by DegU∼P as swarming motility, biofilm formation, SPP1 bacteriophage sensitivity, and exoprotease secretion. In agreement with these observations, transcription of genes regulated by DegU∼P was reduced in the domesticated clone. The transcription of *degU* itself was also reduced (Fig. 5B); since our *degU* transcriptional reporter fusion includes all promoters known to contribute to the expression of the gene (Fig. S3), including the DegU∼P-recognized P3 promoter, it suggests that the I186M substitution affects the auto-regulatory loop that controls the production and activity of DegU (57). Failure to successfully activate the auto-regulatory loop due to the likely impaired binding to DNA of the I186M substitution could be an explanation for the impaired biofilm development, exoprotease production and phage sensitivity, which require high levels of DegU∼P.

The extensive propagation of *B. subtilis* in a nutrient-rich medium, *i.e*., under conditions of relaxed selection for sporulation, resulted in the emergence of a strain that accumulated mutations in genes of biosynthetic pathways, sporulation competence, DNA repair and others (20, 21). Relative to the ancestral, the resulting strain showed different cell and colony morphologies, loss of sporulation and competence, but an overall increased fitness under laboratory conditions (20, 21). Interestingly, our selection, which was also performed in a rich medium, LB, did not result in loss of sporulation. Rather, the I186M substitution modulated the efficiency of sporulation across conditions, specifically, in a host-related condition (RPMI medium) and in LB (Fig. 6). Importantly, the accumulation of DegU was only slightly higher in Ancestral (and *degU^Anc^*) relative to Evolved when sporulation was tested in RPMI, where the sporulation efficiency of Ancestral (and *degU^Anc^*) was also higher than that of Evolved. Since high levels of DegU∼P control production of Spo0A∼P (13), it is possible that I186M places DegU∼P below a threshold required for the stimulation of sporulation through Spo0A. This is consistent with decreased transcription of *degU* by DegU^I186M^ (Fig. 5B). In contrast, no differences in the mean sporulation efficiency of Evolved and Ancestral were found under nutritional conditions (LB medium) that do not support efficient sporulation by most laboratory strains (Fig. 6), and the accumulation of DegU did not differed between Evolved and Ancestral. We note, however, that the accumulation of DegU reflects both transcription/production of the protein and proteolysis; since DegU∼P is a preferred substrate for the ClpXP protease, the steady-state levels of the protein will reflect the ratio of DegU/DegU∼P levels (57). The inability of some domesticated strains of *B. subtilis* to form robust biofilms results from the accumulation of mutations in four chromosomal genes (*sfp*, *epsC*, *swrA,* and *degQ*), in addition to the loss of plasmid-born gene, *rapP* (6). In contrast, under our experimental conditions and using a natural isolate of *B. subtilis*, the target for mutation during domestication is *degU*. Our results are consistent with what was observed in the laboratory strain 168, that has a mutation in *degQ* (6), targeting the DegS-DegU system. Taking into account that targeting the DegS-DegU system is a strategy used by two different strains of *B. subtilis*, with very different genomes, suggests that this system is a major target of adaptation to the laboratory environment.

It seems possible that our experimental conditions did not cause relaxed selection for sporulation; rather, our selection may have first targeted the most costly phenotypes under the test conditions, those directly controlled by DegU∼P, including the formation of complex colonies and robust biofilms, at least in the context of the BSP1 genome. It is interestingly to note that a two-month culture of the Laboratory strain NCIB 3610, resulted in the emergence of strains with different levels of biofilm robustness, as shown by the colony architecture and the expression of genes required for matrix production (3). These phenotypes were the result of mutations in the *sinR* gene, coding for a master regulator of biofilm development, and arose both on plates as well as in LB cultures (3). One conclusion offered was that matrix overproduction can be neutral or advantageous in a rich medium (3). The difference in the mutations obtained under our experimental conditions and the study of Leiman and co-workers is in line with the idea that adaptations to a new environment, depend both on the initial genome as well as the culture history of the strain (80).

Taken together, the results suggest that the I186M mutation impairs the ability of DegU to function as a transcription factor and that this feature confers an advantage to the natural isolate BSP1 when growing under laboratory conditions. Interestingly, *arcA* in *E. coli* and *rpoS* in *S. enterica* and *E. coli*, regulators of stationary phase processes, are also common targets of laboratory adaptation (73, 81). This strengthens the evidence for a general rule that the initial adaptations to a new environment involve changes in genes that act as regulatory hubs of networks that affect the stationary phase of growth.

The accumulation of mutations during adaptation to a laboratory environment over a relatively small number of passages also unraveled a signal of antagonistic pleiotropy: Evolved exhibited changes in traits in host-related environments, and interestingly, it showed increased survival in the presence of macrophages (Fig. 6A). This provides support for the coincidental hypothesis (82, 83) that posits that adaptations to new environments can lead to changes in complex interactions with hosts. Increased survival of the Evolved when facing the cells of the host immune system also implicates DegU potential relevance for the interaction between *B. subtilis* and its host in its natural environment. Importantly, the genus *Bacillus* is commonly used as a probiotic (84, 85) and *B. subtilis* was shown to stimulate macrophage activity and the host immune response (86–89). However, the mechanisms behind *Bacillus* role as a probiotic are still unclear (84). Given that DegU is widely conserved amongst the *Bacillus* genus (Fig. S2), the role of DegU in this interaction should be studied in future work. In addition, our study highlights the importance of performing short-term cultivation of bacterial natural isolates to prevent the loss of traits that may be important for the probiotic activity of *B. subtilis*.

## Material and Methods

### General methods

Lysogeny Broth (LB) medium was used for the routine growth of *B. subtilis* and *Escherichia coli*. The *E. coli* strain DH5α was used as the host strain for the construction and maintenance of plasmids and was grown in the presence of 100 µg ml^-1^ ampicillin when carrying vectors or recombinant plasmids. When appropriate, *B. subtilis* strains were grown in the presence antibiotics, used at the following concentrations: 5 µg ml^-1^ chloramphenicol, 1 µg ml^-1^ erythromycin, 1 µg ml^-1^ neomycin for liquid cultures and 3 µg ml^-1^ neomycin on solid media.

### Other Methods

The construction of all plasmids and strains is described in detail in the Supplemental Material. The *B. subtilis* strains used or constructed in this work are listed in Table S1. Plasmids are listed in Table S2 and oligonucleotides in Table S3. Modeling of the DegU structure is also described in Text S1.

### Domestication experiments

Five independent populations, all derived from the ancestral natural isolate BSP1 (60), were grown for 16 days in LB with a 1:100 dilution into fresh medium every 24 h. This is a common media for growing *B. subtilis* in the laboratory environment and may introduce selective pressure against sporulation and biofilm formation. At the point of dilution, an aliquot from each culture was collected and kept frozen at −80°C for subsequent analysis.

### Whole-genome sequencing

To identify the mutations that emerged after 8 days of evolution we extracted DNA from the Evolved clone from population 1 and from the Ancestral. The DNA library construction and sequencing were carried out by the IGC genomics facility. Each sample was pair-end sequenced on an Illumina MiSeq Benchtop Sequencer. Standard procedures produced data sets of Illumina paired-end 250-bp read pairs. The mean coverage per sample was of 30 and 18, for the Evolved and Ancestral respectively. Mutations were identified using the BRESEQ pipeline version 0.32.1 (90) with default parameters and using the available BSP1 genome (91) as a reference genome. All predicted mutations were manually inspected using IGV (92).

### Macrophages culture and infection assay

The murine macrophage cell line RAW 264.7 was cultured in RPMI medium (Sigma), supplemented with 2 mM L-glutamine, 1 mM sodium pyruvate, 10 mM hepes, 50 µM 2-mercaptoethanol solution and 10 % heat-inactivated Fetal Bovine Serum in an atmosphere of 5 % CO2. For the infection, *B. subtilis* and the macrophages were grown separately in a 24-well tissue plate containing fresh RPMI media as described above. At 24h of acclimatization, *B. subtilis* was diluted 1:100 into fresh RPMI. Macrophages were washed, re-suspended in fresh RPMI and activated with 2 µg ml-1 CpG for another 24h (93). Then, the macrophages were washed to remove the remaining CpG, fresh RPMI media was added and *B. subtilis* added to a 1:8 MOI (multiplicity of infection; about 8 x 10^6^ cells). At the indicated time points of infection, the wells were scrapped and the contents centrifuged at 6000 g for 10 minutes at room temperature. After centrifugation, the samples were serially diluted and plated to determine the titer of total, viable, cells and heat-resistant spores.

### Sporulation assays

Sporulation of *B. subtilis* was usually analyzed in LB and in supplemented RPMI. When using LB, *B. subtilis* cultures were grown overnight, diluted 1:100 and incubated for 24 h at 37 °C. At this time, dilutions of the cultures were plated for total viable counts and treated for 20 min at 80°C to determine the titer of heat-resistant spores. For supplemented RPMI, the cultures were grown as described above for 48h and plated for viable cells and spore counts as described above for LB. The sporulation efficiency was defined as the ratio of heat-resistant spores relative to the total viable cell count (18).

### SPP1 phage lysates and transduction

SPP1 lysates were prepared as described by Yasbin and Young (94). Briefly, a dense culture of *B. subtilis* was infected with different dilutions of SPP1 in a semisolid LB agar (LB containing 0.7% agar). The plate containing near confluent phage plaques was washed with 4 ml of TBT, centrifuged at 5000 g for 10 min, treated with 12 µg ml^-1^ DNase and filtered through a 0.45 µm syringe filter. The indicator strain PY79 was used for titration of the SPP1 lysates as described by São-José et al (95). SPP1 phage transduction was performed as described (42). The recipient strains were grown in LB until stationary phase after which 1 ml of the culture of the recipient strain was mixed in a glass tube with 10 mM CaCl_2_ and infected with an MOI of 1 of the donor SPP1 lysate. The transduction mixture was then incubated at 37°C for 25 min with agitation, centrifuged at 5000 g for 10 min, washed with 2 ml of LB, and centrifuged again at 5000 g for 10 min. The supernatant was discarded, the pellet was resuspended in 100 µl of LB and plated onto LB plates fortified with 1.5 % agar with the appropriate antibiotics and 10 mM of sodium citrate.

### Competence assay

Development of competence was performed as described by Baptista et al (47). Briefly, *B. subtilis* cultures were grown overnight and diluted 1:100 in GM1 at 37 °C. Ninety minutes after the end of the exponential growth, the cultures were diluted 1:10 in GM2 and incubated for 90 minutes at 37 °C. At this point, a sample of the cultures was serially diluted in LB and plated for determination of total colony forming units (CFU) per milliliter. For transformation, DNA from strain AH7605 or W648 was added to 500 µl of the culture samples, to a concentration of 5 µg ml^-1^, the mixture incubated for 30 min at 37 °C and finally plated with the appropriate antibiotics. The transformation efficiency is the ratio between the number of transformants and the total number of colonies.

### Protease activity assay

Secreted proteases were observed essentially as described by Saran et al (96). The strains were grown until they reached an absorbance of 0.8 at 600 nm. At this time the cultures were diluted to an absorbance of 0.01 at 600 nm. 10 µl of this dilution was spotted in a 2 % skimmed milk plate and incubated at 37 °C for 48 h. Then, 6 ml of 10 % Tannic Acid was added for detection of the protease-positive strains. The diameter of the halos observed was measured, and the diameter of the colony was subtracted in order to obtain the real value of the halo.

### Swarming and colony morphology assays

Swarming motility was examined according to the method described by Kearns and Losick (42). For colony morphology, the *B. subtilis* cultures were grown overnight and 3 µl of the culture was spotted onto an MSgg (97) plate fortified with 1.5 % agar. The plates were incubated at 28 °C or 37 °C. The images were captured at the times indicated in the figures.

### Biofilm quantification by crystal violet

The method used for estimating the solid-surface-associated biofilm formation with crystal violet was as described by Morikawa et al (98). Briefly, an overnight culture was diluted to an absorbance of 0.03 at 600 nm and mixed 1:100 into 100 µl of MSgg in a 96-well plastic titer plate. The plate was incubated for 48 h at 25 °C. Then, the culture was carefully removed from the wells. After washing two times with distilled water, 150 µl of 1 % crystal violet was added to the wells and incubated for 25 min at room temperature. The wells were washed again two times with distilled water and the crystal violet attached to the biofilm matrix was solubilized in 150 µl of DMSO and incubated for 10 minutes at room temperature. The removed culture was quantified by measuring its absorbance at 600 nm and the biofilm attached to the crystal violet was quantified measuring its absorbance at 570 nm.

### Biofilm fluorescence imaging

For biofilm imaging, the *B. subtilis* cultures were grown overnight and 3 µl of the culture was spotted onto an MSgg plate fortified with 1.5 % agar and incubated for 96h at 28°C. Images were acquired on a Zeiss Axio Zoom.V16 stereomicroscope equipped with a Zeiss Axiocam 503 mono CCD camera and controlled with the Zeiss Zen 2.1 (blue edition) software, using the 1x 0.25 NA objective, the fluorescence filter set GFP and the Bright Field optics.

### Fluorescence microscopy and image analysis

Cultures were grown in LB until one hour after the end of the exponential phase. The cells were collected by centrifugation (1 min at 2.400 x g, room temperature), and washed with 1 ml of phosphate-buffered saline (PBS). Finally, the cells were resuspended in 100 μl of PBS and applied to microscopy slides coated with a film of 1.7% agarose. Images were taken with standard phase contrast and GFP filter, using a Leica DM 6000B microscope equipped with an aniXon+EM camera (Andor Technologies) and driven by Metamorph software (Meta Imaging series 7.7, Molecular Devices). For quantification of the GFP signal, 6×6 pixel regions were defined in the desired cell and the average pixel intensity was calculated and corrected by subtracting the average pixel intensity of the background, using Metamorph software (Meta Imaging series 7.7, Molecular Devices).

### Immunoblot analysis

Immunoblot of DegU was analyzed in LB and in supplemented RPMI. When using LB, *B. subtilis* cultures were grown until one hour after the end of the exponential phase and samples (10 ml) were withdrawn. For supplemented RPMI, the cultures were grown as described above and samples (10 ml) were withdrawn. In both media, the cells were collected by centrifugation (5 min at 15300 x g, 4° C). The cells were resuspended in 1ml Lysis buffer (50 mM NaH_2_PO_4_, 0.5 M NaCl, 10 mM Imidazole, pH 8.0) and whole-cell lysates prepared using a French press cell (19,000 lb/in^2^). Proteins in the lysates (10 µg) were then separated on 15% SDS-PAGE gels and the gels subject to immunoblot analysis using an anti-DegU antibody of established specificity at a 1:1000 dilution (99). Gels run in parallel were stained with Coomassie brilliant blue to be used as loading controls.

## Data availability

Genome sequencing data have been deposited with links to BioProject accession number PRJNA592868 in the NCBI BioProject database (https://www.ncbi.nlm.nih.gov/bioproject/).

## Acknowledgments

We thank Dr. Mitsuo Ogura for the gift of the anti-DegU antibody, Dr. Kazuo Kobayahis for the gift of strains WTF28 and W648, Dr. Jan-Willem Veening for the gift of strains 08G52, 08G57, and 08I09, and Dr. Carlos São José for the gift of the SPP1 lysate and advice on its use. We thank Tanja Ðapa for critically reading this manuscript. We also thank Mónica Serrano for helpful discussions.

This work was supported by Project LISBOA-01-0145-FEDER-007660 (“Microbiologia Molecular, Estrutural e Celular”) funded by FEDER funds through COMPETE2020 –“Programa Operacional Competitividade e Internacionalização” (POCI), by national funds through the FCT (“Fundação para a Ciência e a Tecnologia”) grant POCI/BIABCM/60855/2004 to A.O.H. and partially supported by PPBI - Portuguese Platform of BioImaging (PPBI-POCI-01-0145-FEDER-022122) co-funded by national funds from OE - “Orçamento de Estado” and by European funds from FEDER. The research leading to these results also received funding from the European Research Council under the European Community’s Seventh Framework Programme (FP7/2007–2013)/ERC grant agreement no 260421 – ECOADAPT to I.G. H.C.B. was the recipient of a doctoral fellowship from the FCT (PD/BD/128429/2017). The funders had no role in study design, data collection and analysis, decision to publish, or preparation of the manuscript.

