## Supplemental Material for "Rampant loss of social traits during domestication of a *Bacillus subtilis* natural isolate"

**Short title:** *Domestication of a Bacillus subtilis natural isolate*

**Keywords:** DegU; experimental evolution; biofilm development; sporulation; extracellular proteases.

### Text S1 Supporting information text

### Supporting Material and Methods

**Plasmid and strain construction.** Strain HB4 is a derivative of the wild-type strain BSP1 (1), obtained by moving the *degU::cat* mutation present in strain WTF28 (2) using SPP1-mediated generalized transduction. The *degU::cat* mutation in HB4 was then complemented as follows (Fig. S2): *i*) for complementation with *degU*$^{Anc}$, the promotor region of the *degSU* operon and the *degU* gene were PCR amplified using primer pair DegS-438F/DegS+64R and DegS+927F/DegU+775R respectively. An overlapping PCR using primers DegS-438F and DegU+775R generated a fragment which was digested with HindIII and BamHI; *ii*) for complementation with *degU*$^{Evo}$, two PCR products that included the I186M mutation in *degU* were obtained with the primer pairs DegS+927F/DegU+555R and DegU+555F/DegU+775R and an overlapping PCR was obtained from the two fragments. A final overlapping PCR fragment was obtained with primers DegS-438F and DegU+775T, which was digested with HindIII and BamHI. All the final overlapping PCR products were then inserted between the HindIII and BamHI sites of pMLK83, an *amyE* integrational vector carrying a neomycin-resistance determinant selectable in *B. subtilis* in a single copy (3). The resulting plasmids, pHB7 (carrying *degU*$^{A10E}$) and pHB8 (carrying *degU*$^{Evo}$) were used to transform *E. coli* DH5α. Note that since *degU*$^{Anc}$ is toxic in *E. coli* (4) and *degU*$^{Evo}$ is not, pHB7 was designed to carry a mutation in degU that codes for a form of the protein with the single amino acid substitution A10E, which is not toxic for *E. coli*. The resultant recombinant plasmids, pHB7 (carrying *degU*$^{A10E}$) and pHB8 (carrying *degU*$^{Evo}$) were then used to transform BSP1 by co-transformation with genomic DNA of strain HB4 to produce strains HB7 and HB8 (Fig.

S2). For changing the chloramphenicol resistance determinant to an erythromycin resistance determinant in HB7 and HB8, both strains were transformed with pCM::ery originating the resulting erythromycin-resistant strains HB11 and HB12, respectively (Fig. S2). The loss of chloramphenicol resistance was verified in HB11 and HB12. To correct the *degU*$^{A10E}$ mutation, the fragment obtained by overlapping PCR in the construction of pHB8 was digested with EcoRI and the *amyE* integrational vector pDG364 was digested with BamHI. Then, both the PCR fragment and the plasmid were treated in a first step with the Klenow fragment of DNA polymerase to originate blunt ends and afterwards digested with HindIII. The resultant fragment and plasmid were then ligated and used to transform *E. coli* DH5α to produce pHB1. Transformation of HB11 with pHB1 produced strain HB13 (Fig. S2). Transformation of HB12 with pHB1 produced strain HB14 (Fig. S2). Note that plasmids pHB1, pHB7 and pHB8 were sequenced with primers DegS-438F and DegU+775R to verify the presence of the desired sequences and the absence of unwanted mutations. For the construction of the $P_{aprE}$-, $P_{hag}$-, and $P_{degU}$-*gfp* fusions we extracted the DNA from the strains 08G57 ($P_{aprE}$-*gfp*), 08G52 ($P_{hag}$-*gfp*), and 08I09 ($P_{degU}$-*gfp*). The extracted DNA was then used to transform the Ancestral, Evolved and Lab strains. For the construction of $P_{degU}$-*gfp* fusions, we transformed the Ancestral, Evolved and Lab strains with the plasmid pIP11. Development of competence and transformation was performed as described by Yasbin et al (5). Transformants were then selected for their appropriate antibiotic resistance and confirmed by fluorescence microscopy.

**Modeling of the DegU structure**. The model for full-length DegU was built using the crystal structure of the beryllofluoride-activated LiaR protein from *Enterococcus faecium* (PDB code: 5hev) as the model (6). The DegU model was generated by comparative

modeling with Rosetta (7) using evolutionary coupling-derived distance restraints (8). The DNA-binding domain of DegU was independently modeled using as the model
**M**KGNIVQYNFADIEEEVYSLDYAIAWNTNEENVNIIPFTNKFCKESIESFCLGKINNFVEI
LNEGFVENHHYVHLDK**M**ISVPKKKVNLVYQQDTHGYLLRDDNDNLIPAKITSEQSKSIS
SK**M**ELFCAGEEKCLINILLKADPSYILDVDSIKDKNILNLGYESIDRYKEYNFDDDKILIFFI
NKKRYSVI**M**KKTNNSDNDLVSRNNAIKELFTNKAGNLN template the crystal structure of the wild type DNA binding domain from *E. faecalis* LiaR complexed with a 22bp DNA fragment (PDB code: 4wuh) (9).

### Supporting Results and Discussion

#### The DegU model

DegU belongs to the NarL/FixJ family of transcription factors which have a helix-turn-helix (HTH) motif of about 65 amino acid residues close to their C-terminus (10). Using the HHpred server (11) we have identified the structure of activated transcriptional regulatory protein LiaR (PDB: 5HEV) from *Enterococcus* faecium, also a member of the NarL/FixJ family, as the structural homolog with the highest sequence identity/similarity (37%/63.4%) to *B. subtilis* DegU. LiaR is a regulator of cell envelope stress in many Gram-positive bacteria, including *B. subtilis*, and is phosphorylated by a membrane-bound histidine kinase, LiaS (12). As other members of the family, LiaR *of E. faecium* consists of two functional domains, a receiver domain (residues 1-139 of the 206-residues long protein) which is the site of phosphorylation, at a conserved aspartate, by LiaS, and a DNA-binding domain (residues 140-210), which bears a HTH motif (6, 9).

Based on the structure of LiaR, DegU is modelled as a homodimer composed of two conserved domains (7, 8): an N-terminal receiver domain (RD, residues 1-120 of the 229-residues long protein) connected by a linker to a C-terminal DNA-binding domain (DBD, residues 160-225), bearing a helix-turn-helix (HTH) motif (residues 182-207) (Fig. 2A and Fig. S3). The HTH motif in DegU is formed by helices 8 (the DNA recognition helix) and 9 (the scaffolding helix) (see Fig. 2B). Ramachandran plots (not shown) reveal that 359 residues (or 89.8%) are located in most favored regions, 36 residues (9%) in additional allowed regions, 1 residue (0.2%) in generously allowed regions, and only 4 residues (or 4%) in disallowed regions. Thus, the model is of high quality. For the DBD, we also created a model of its DNA-bound form using as the crystal structure of LiaR DBD complexed with DNA as the template (9).

**The effects of the I186M, H200Y and V131D substitutions in DegU**

Previous studies have shown that single alanine substitutions of I186 (in the DNA recognition helix 8 of the HTH motif) or H200 (in the scaffolding helix 9), strongly impaired *comG-lacZ* expression, as a direct indicator of *comK* transcription, or *aprE-lacZ* expression (13). Moreover, the H200A substitution impaired binding of purified DegU to the *comK* and *aprE* promoters (13). Furthermore, electrophoretic mobility shift assays also suggested that DegU formed multimeric complexes at both promoters (13).

Residue I186 is conserved in NarL, ComA, and LuxR; as shown in the crystal structure of a NarL-DNA complex, and suggested by our model, I186 contacts bases in the major groove of DNA (10, 13, 14) (Fig. 2B). H200, on the other hand, which is conserved in NarL, contacts the sugar-phosphate backbone of DNA and this is also suggested by our model of a DegU-DNA complex (13) (Fig. 2A). Thus, both the I186M and H200Y substitutions are likely to affect the binding of DegU to its target sequences in DegU-responsive promoters. This inference, in turn, is in line with the strong reduction in expression of the DegU-responsive *gfp* promoter fusions in Evolved (Fig. 5) and the phenotypes associated with this strain (Fig. 3-6, Fig. S4 and S5).

On the other hand, residue V131 is located in a patch of hydrophobic amino acids, conserved among DegU homologs, located just downstream of the end of the Receiver Domain (Fig. S3). This patch is conserved in *E. faecium* LiaR, in which the equivalent residue is V125 (Fig. S3). In the unphosphorylated form of LiaR, as in other members of the NarL/FixJ family such as *Staphylococcus aureus* VraR, the DNA-binding domain is packed against the receiver domain, and the interdomain interaction is stabilized by the linker between the two domains (9, 15). Phosphorylation induces a conformational change in the receiver domain, accompanied by a reorientation of the linker, that releases the DNA-binding domain (9, 15). This allows dimerization of the protein but evidence also suggests a monomer-dimer-tetramer equilibria (9, 15). Higher-order multimerization of LiaR and VraR has been suggested to allow the protein to bind to a range of promoters, which have different arrangements (direct or inverted repeats) and copies of their cognate binding sites (9, 15). As for LiaR, the number and orientation of DegU binding sites in its target promoters vary greatly, and evidence suggests both dimerization and tetramerization of DegU upon phosphorylation (16). Since V131D introduces a negative charge in the conserved hydrophobic pocket at the beginning of the interdomain linker, we speculate that this substitution may affect the ability of DegU to form multimers, or otherwise the orientation of the linker between the Receiver and DNA-binding domains, and in either case its ability to bind to DNA at different promoters.

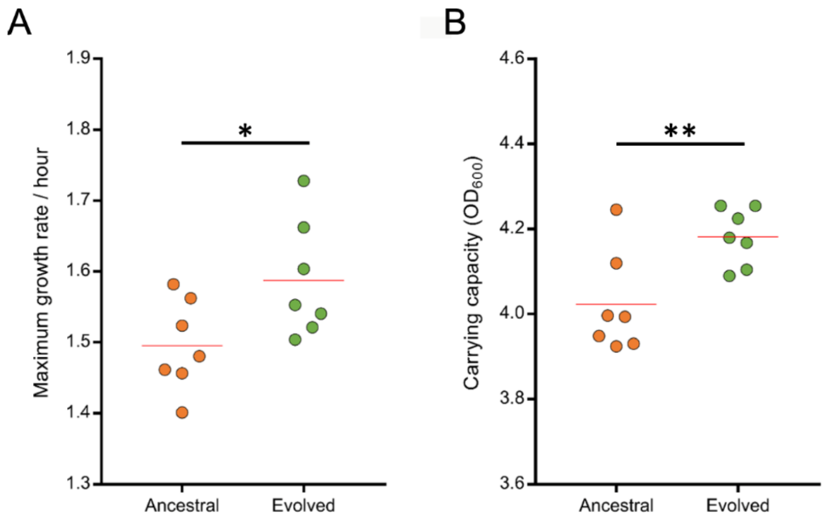


**FIG S1 Evolved has increased growth traits than Ancestral.** (A) Comparison of the maximum growth rate of Ancestral (n = 7) and Evolved (n = 7) in LB, obtained with the R package *growthrates*. *p = 0.04 (B) Comparison of the carrying capacity of Ancestral (n = 7) and Evolved (n = 7) in LB. **p = 0.01. In both panels the Unpaired t test was used. The red line represent the mean.

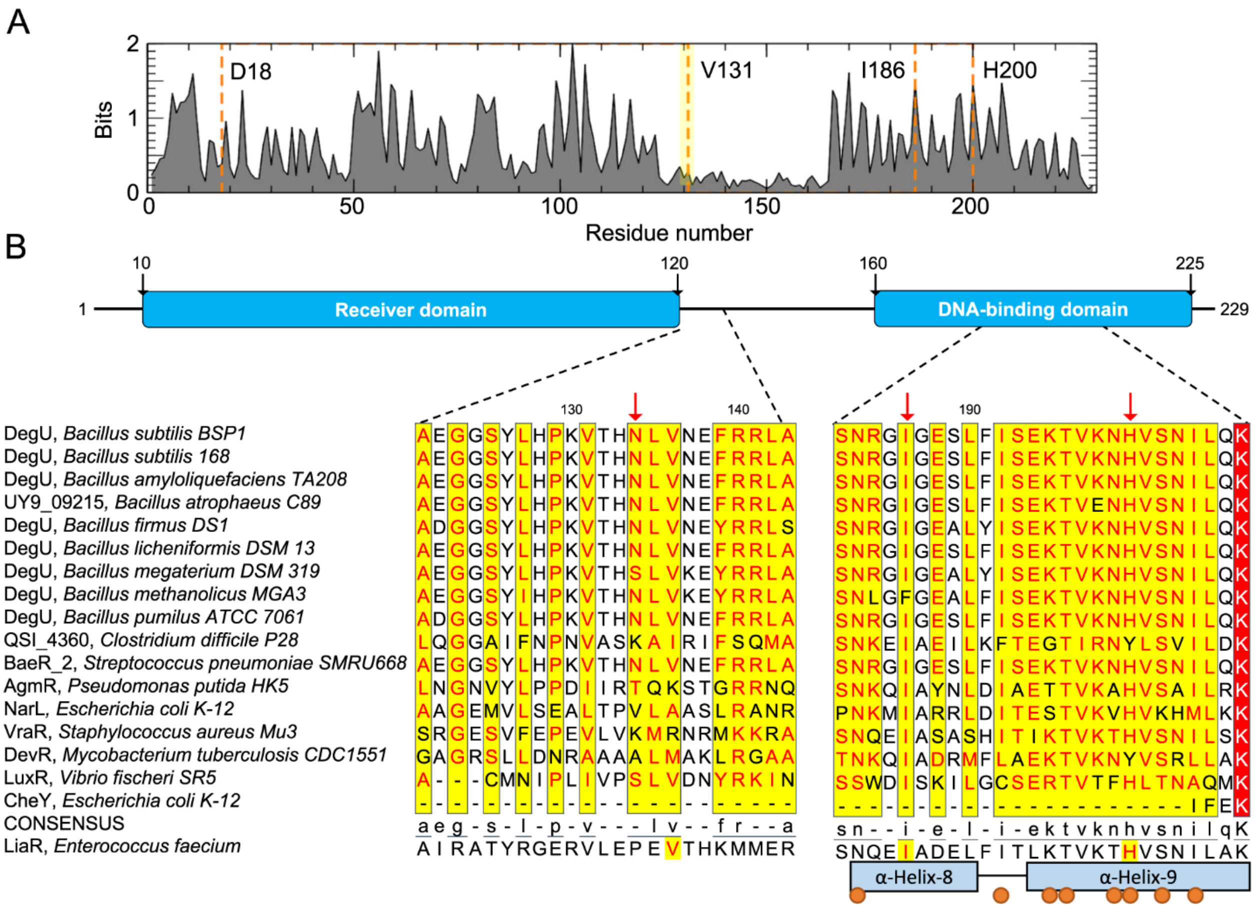


**FIG S2 Alignment of DegU orthologs.** (A) Sequence conservation profile of DegU homologs computed from a Hidden Markov Model multiple sequence alignment using Skylign (17). The overall height indicates the conservation per position. The orange dashed line shows the position of D18, V131, I186, and H200. With the exception of V131, all other residues are in highly conserved regions; V131, however, is located in a conserved hydrophobic patch (expanded in panel B). (B) Alignment of the regions close to the observed substitutions in DegU. The panel was produced with ESPript 3.0 (18) following a Clustal Omega (19) alignment of the selected sequences with the accession codes AGA21795, NP_391429, AEB25746, EIM10937, EWG11477, AAU25236, ADF41907, AIE61365, EDW22531, EQJ51456, COC86792, A8R3T0, P0AF28, P0C0Z1,

P9WMF8, EHN68297, and P0AE67. The red arrows indicate the residues which are the site of the V131D, I186M, and H200Y substitutions herein described. The boxes indicate blocks of high sequence identity; red indicates chemical similarity. The sequence of *E. faecium* LiaR (accession code EPI11259), used for the homology modeling of the *B. subtilis* DegU protein, is indicated below the consensus. The positions of helices 8 (scaffolding helix) and 9 (DNA recognition helix) is also shown. The brown dots indicate residues important for binding of DegU to the *comK* and *aprE* promoters (13).

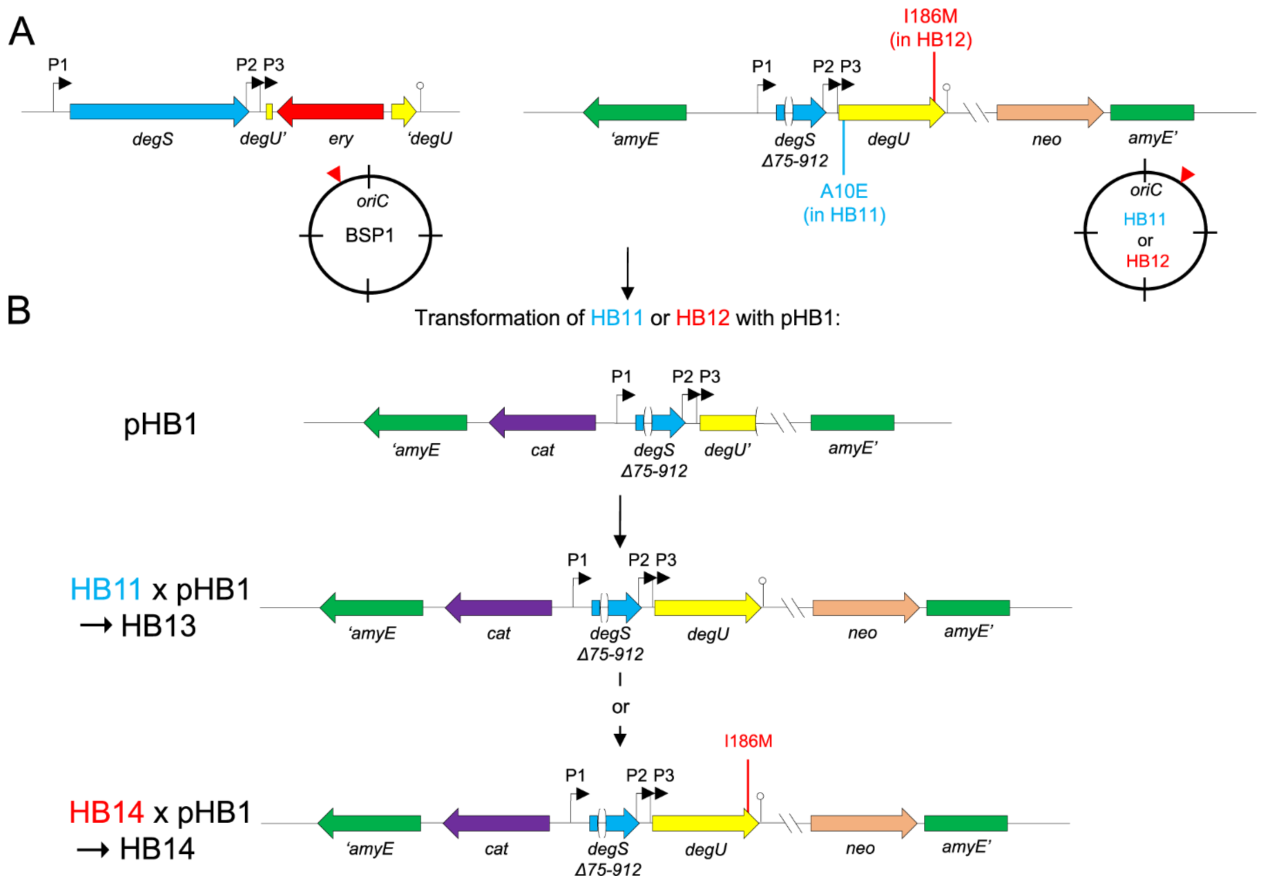


**FIG S3 Construction of strains bearing *degU*$^{Anc}$ and *degU*$^{Evo}$ at an ectopic site.** The figure depicts the construction of strains HB13 and HB14. (A) left: genome organization of the native *degU* locus, at the left of *oriC,* as shown in the circle below the genetic map, in Ancestral. Note the presence of the three promoters, P1 to P3, that drive expression of *degU.* Right: strains HB11 and HB12 are BSP1 derivatives bearing a *degU*::*em* insertion at the *degU* normal locus and an insertion of the *degU* region at the non-essential *amyE* locus, to the right of *oriC*, as depicted. The region inserted at *amyE* includes a *degS* in frame-deletion that removes nucleotides 75-912 of the coding region, so that the strain has only one copy of the gene, at the normal locus. The *degU* allele inserted at *amyE* codes for a form of the protein with the A10E substitution, in strain HB11 (blue), or for the

I186M substitution, in strain HB12 (red), as shown. Note that in both strains, expression of *degU* from *amyE* can still occur from P1 to P3. (B) The panel depicts the result of transforming HB11 or HB12 with plasmid pHB1. Using HB11 as the recipient, a wild-type degU allele is restored, yielding HB13. Using HB12 as the recipient, the mutation leading to the A10E substitution is corrected, yielding strain HB14, which expresses $degU^{Evo}$ ($degU^{I186M}$) from *amyE* (see S1 Text for details).

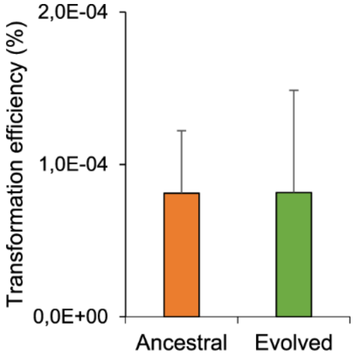


**FIG S4 Competence is not affected in Evolved.** Transformation of Ancestral and Evolved with genomic DNA from AH7605 (Ancestral, n = 3, Evolved, n = 3). The transformation efficiency is expressed as the ratio between the number of transformants obtained and the total number of colonies. The Unpaired t test was used. $p = 0.99$. The error bars represent the standard deviation.

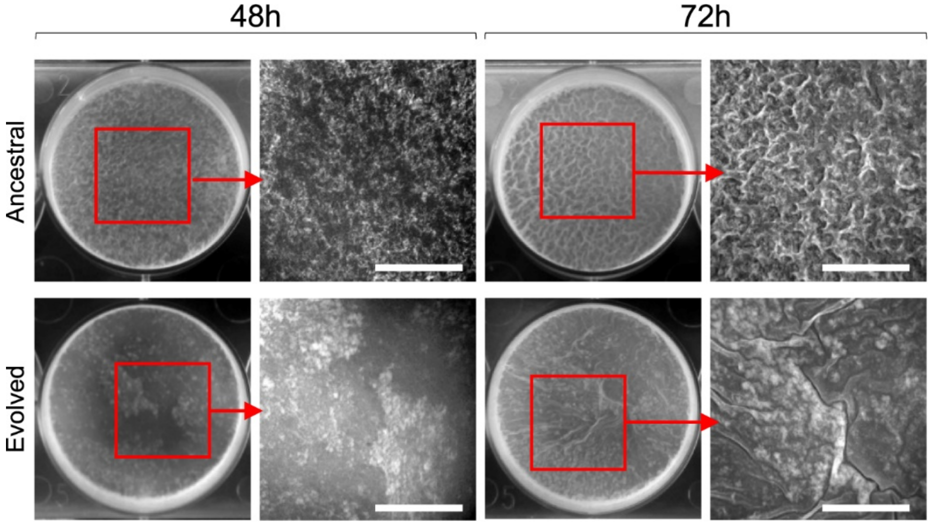


**FIG S5 Pellicle formation is impaired in Evolved.** Representative images of pellicle formation by Ancestral and Evolved after incubation in liquid MSgg medium at 28ºC for the indicated time, in hours. The region boxed in red in the two sets of panels on the left, for each time sample, are magnified on the right. Scale bar, 1 cm.

**TABLE S1** Bacterial strains used in this study.

| Strain | Relevant genotype/phenotype[a] | Origin/Construction |
|---|---|---|
| PY79[b] | Prototrophic | Laboratory stock |
| MB24 | *trpC2 metC3*/Spo$^+$ | “ |
| JH642 | *trpC2 pheA1*/ Spo$^+$ | “ |
| 168 | Prototrophic | “ |
| BSP1[c] | Gastrointestinal isolate #200 / Prototrophic | (20) |
| B081#1[d] | Clone isolated from population 1 after 8 days of evolution | This work |
| WTF28 | *degU::cat* /Cm$^R$ | (2) |
| 08G52 | 168 derivative*, hag-gfp,* Cm$^R$ | (21) |
| 08G57 | 168 derivative, *aprE-gfp*, Cm$^R$ | (21) |
| 08I09 | 168 derivative*, amyE::PdegSU-gfp,* Sp$^R$ | (21) |
| HB4 | BSP1 derivative, *degU::cat*, Cm$^R$ | This work |
| HB7 | BSP1 derivative, *degU::cat amyE:: degU$^{A10E}$*, Cm$^R$ Neo$^R$ | “ |
| HB8 | BSP1 derivative, *degU::cat amyE::degU$^{Evo}$*, Cm$^R$ Neo$^R$ | “ |
| HB11 | BSP1 derivative, *degU::ery amyE:: degU$^{A10E}$*, Em$^R$ Neo$^R$ | “ |
| HB12 | BSP1 derivative, *degU::ery amyE::degU$^{Evo}$*, Em$^R$ Neo$^R$ | “ |
| HB13 | BSP1 derivative, *degU::ery amyE::degU$^{Anc}$*, Cm$^R$ Neo$^R$ Em$^R$ | “ |
| HB14 | BSP1 derivative, *degU::ery amyE::degU$^{Evo}$*, Cm$^R$ Neo$^R$ Em$^R$ | “ |
| HB33 | BSP1 derivative, *aprE-gfp*, Cm$^R$ | “ |
| HB34 | BSP1 derivative, *hag-gfp*, Cm$^R$ | “ |
| HB35 | BSP1 derivative, *amyE::P$_{degSU}$-gfp*, Sp$^R$ | “ |
| HB36 | B081#1 derivative, *aprE-gfp*, Cm$^R$ | “ |
| HB37 | B081#1 derivative, *hag-gfp*, Cm$^R$ | “ |
| HB38 | B081#1 derivative, *amyE::P$_{degSU}$-gfp*, Sp$^R$ | “ |
| HB39 | PY79 derivative, *aprE-gfp*, Cm$^R$ | “ |
| HB40 | PY79 derivative, hag-*gfp*, Cm$^R$ | “ |
| HB41 | PY79 derivative, *amyE::P$_{degSU}$-gfp*, Sp$^R$ | “ |
| HB42 | BSP1 derivative, *amyE::P$_{bslA}$-gfp*, Cm$^R$ | “ |
| HB43 | B081#1 derivative, *amyE::P$_{bslA}$-gfp*, Cm$^R$ | “ |

| | | |
|---|---|---|
| HB44 | PY79 derivative, *amyE::P*$_{bslA}$*-gfp*, Cm$^R$ | " |

[a]Cm$^R$, chloramphenicol resistance; Neo$^R$, neomycin resistance; Em$^R$, erythromycin resistance; Sp$^R$, spectinomycin resistance. [b]Herein termed "Lab". [c]Herein termed "Ancestral". [d]Herein termed "Evolved".

**TABLE S2** Plasmids used in this study.

| Plasmid | Relevant genotype/Phenotype | Origin |
|---|---|---|
| pMLK83 | Integration vector, allows ectopic integration at *amyE*; $Amp^R$ $Neo^R$ | (3) |
| pDG364 | Integration vector, allows ectopic integration at *amyE* locus; $Amp^R$ $Neo^R$ | BGSC[a] |
| pCM::ery | For the replacement of the chloramphenicol resistance marker by an erythromycin resistance determinant | BGSC[a] |
| pHB1 | pDG364 derivative carrying $degU_{EcoRI}$ | This work |
| pHB7 | pMLK83 derivative carrying $degU^{A10E}$ | “ |
| pHB8 | pMLK83 derivative carrying $degU^{Evo}$ | “ |
| pIP11 | pMLK83 derivative carrying $P_{bslA}$-*gfp* | Laboratory stock |

[a]*Bacillus* Genetic Stock Center.

**TABLE S3** Oligonucleotide primers used in this study.

| Name | Sequence (5' to 3')[a] |
|---|---|
| DegS-438F | TGTAAAGCTTGGTTCCCCGTC |
| DegU+775R | GCTTGGATCCCTGCCTTATTG |
| DegS+64R | AGATTCAGAATGCTTTAGCGCGCTCCCGTCAACGGTTTTCAG |
| DegS+927F | GCGCTAAAGCATTCTGAATCTGAAGAAATT |

[a]Underlined sequences represent introduced restriction sites.
